# A Front-Back Confusion Metric in Horizontal Sound Localization: The FBC Score

**DOI:** 10.1101/2020.02.12.945303

**Authors:** Tim Fischer, Marco Caversaccio, Wilhelm Wimmer

**Affiliations:** Hearing Research Laboratory, ARTORG Center for Biomedical Engineering Research, University of Bern, Bern 3008, Switzerland and the Department of ENT, Head and Neck Surgery, Inselspital, Bern University Hospital, University of Bern, Bern 3008, Switzerland

**Keywords:** Binaural cues, cone of confusion, cochlear implants, diagnostics in audiology, spatial hearing outcome measures

## Abstract

In sound localization experiments, currently used metrics for front-back confusion (FBC) analysis weight the occurring FBCs equally, regardless of their deviation from the cone of confusion. To overcome this limitation, we introduce the FBC Score. A sound localization experiment in the horizontal plane with 12 bilaterally implanted cochlear implants (CI) users and 12 normal hearing subjects was performed to validate the method with real clinical data. The overall FBC Rate of the CI users was twice as high as the FBC Score. For the control group, the FBC Rate was 4 times higher than the FBC Score. The results indicate that the FBC Rate is inflated by FBCs that show a considerable deviation from the corresponding value on the cone of confusion.

## I. Introduction

SOUND source localization primarily relies on the evaluation of interaural time and level differences. Front-back confusions (FBCs) occur because sound sources on the cone of confusion are equidistant from the left ear and the right ear and thus provide identical interaural time and level differences for a listener [1]. An illustration for a sound source on the cone of confusion is given in Figure 1.

**Fig. 1.**
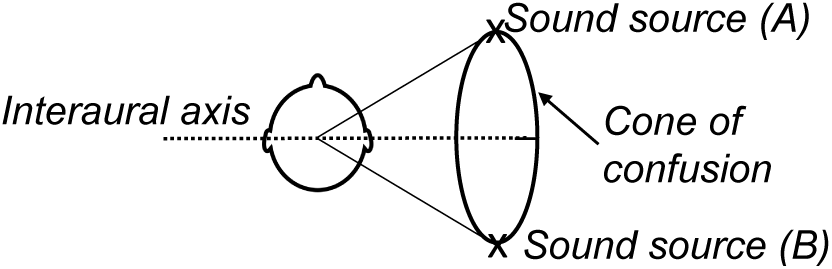
The cone of confusion is defined as an imaginary cone extending outward from the center of the head [2]. The axis of the cone is defined by the interaural axis. Due to the symmetry of this cone around the interaural axis, sound source (A) and (B) produce identical interaural time and level differences as they are equidistant from the left ear and the right ear. Sound sources on the cone of confusion are thus useless for binaural sound localization.

FBC errors are not simply outliers with extreme error amplitudes. The underlying cause is different from normal localization errors (see Figure 1) which is why FBCs should be analyzed seperately [3]–[7].

There is a continuing debate in the literature on the definition of an FBC error. In horizontal sound localization experiments, the most common definition classifies responses crossing the interaural axis as FBCs [3]–[6], [8], [9]. This definition, which defines the FBC Rate, is sufficient for experiments that only require a coarse angular resolution of the test setup or feedback method [7], [10], [11]. However, such setups limit the measurement resolution for sound localization accuracy.

A refined FBC definition requires FBC responses to fall within a specific range of a response-dependent regression line, which is mirrored on the interaural axis [11], [12].

Further definitions as to what constitutes a FBC error are defined by fixed threshold values. These thresholds refer either to a minimum required deviation of the given response in relation to the interaural axis or to a maximum allowed deviation between the response and the stimulus mirrored on the interaural axis [13]. There is no general consensus regarding the magnitude of the thresholds.

Apart from thresholds that are not standardized, the current limitation of FBC analysis approaches is that all FBCs are considered equally strong. The deviation from the measurement position to which the error refers, which is the deviation of the response from the stimulus position mirrored on the interaural axis, is not considered. In addition, special care should be taken with stimuli presenting close to the interaural axis, as localization errors overlap with FBC errors in this area.

Herein, we propose a metric that allows to quantify the impact of FBCs on sound localization outcomes in the horizontal plane more precise than with the commonly used FBC Rate. The FBC Score can be applied regardless of the number of available localization results, the localization performance of the subjects and the measurement setup.

## II. Materials and Methods

### A. Study design and participants

This study was designed in accordance with the Declaration of Helsinki and was approved by the local institutional review board (KEK-BE, No. 2018-00901). Twelve bilaterally implanted cochlear implants (CIs) users participated in the study, all using Sonnet (Med-El GmbH, Innsbruck, Austria) processors with an omnidirectional microphone setting. The CI users had a monosyllabic word recognition score in quiet of 70% or better at 60/65 dB_SPL_. For comparison, a control group of 12 normal hearing (NH) adults was included.

### B. Static Sound Source Localization Test

Static sound source localization was performed with 12 equally spaced loudspeakers arranged in a circle around the subject. The test stimulus consisted of pink noise with 200 ms length. To prevent the use of monaural level cues, level roving between 60 to 70 dB_SPL_ was applied. In total, 36 stimuli per subject were played, 3 stimuli per loudspeaker. The order of the stimuli with respect to the loudspeaker was randomized [10]. The loudspeakers were hidden behind a sound transparent curtain, no prior knowledge about possible stimuli directions was provided to the subjects. The subjects’ feedback on the perceived location of the stimulus was recorded via a graphical user interface showing a dial with a resolution scale of 1-degree angle and a login button.

### C. Calculation of the FBC Score

We define the outcome of a sound localization experiment as a set of stimuli 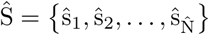 and corresponding subject responses 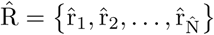, where 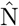 denotes the total number of test items. In relation to the FBC Rate (see Eq. 1), only responses that do cross the interaural axis are considered for the FBC analysis. After exclusion, we obtain a reduced set of stimuli S = {s_1_, s_2_, …, s_N_} and responses R = {r_1_, r_2_, …, r_N_} with N items.

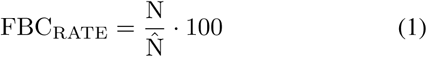

The ideal position of an FBC, i.e., the position of a stimulus s_i_ mirrored on the interaural axis, is defined as the FBC center c_i_. We compute the deviation *θ*_*i*_ between the response r_i_ and the FBC center c_i_ as the shortest absolute angular difference (minor arc). An example for one stimulus-response pair is given in Figure 2. The maximum allowable deviation *θ*_max,i_ is defined by the interaural axis and the FBC center c_i_ (see Figure 3). The farther away a response r_i_ is from its corresponding FBC center and the closer to the interaural axis, the less likely it is to be an FBC. Therefore, we introduce a weighting factor w_i_ for each measured FBC:

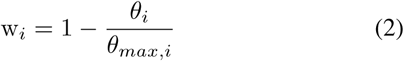

where *θ*_*max,i*_ is the maximum deviation in the direction of the response (clockwise or counterclockwise) under consideration of the interaural axis. The weighting w_*i*_ ranges from 0 (response on the interaural axis) to 1 (a perfect FBC). An illustration of the procedure is given in Figure 3.

**Fig. 2.**
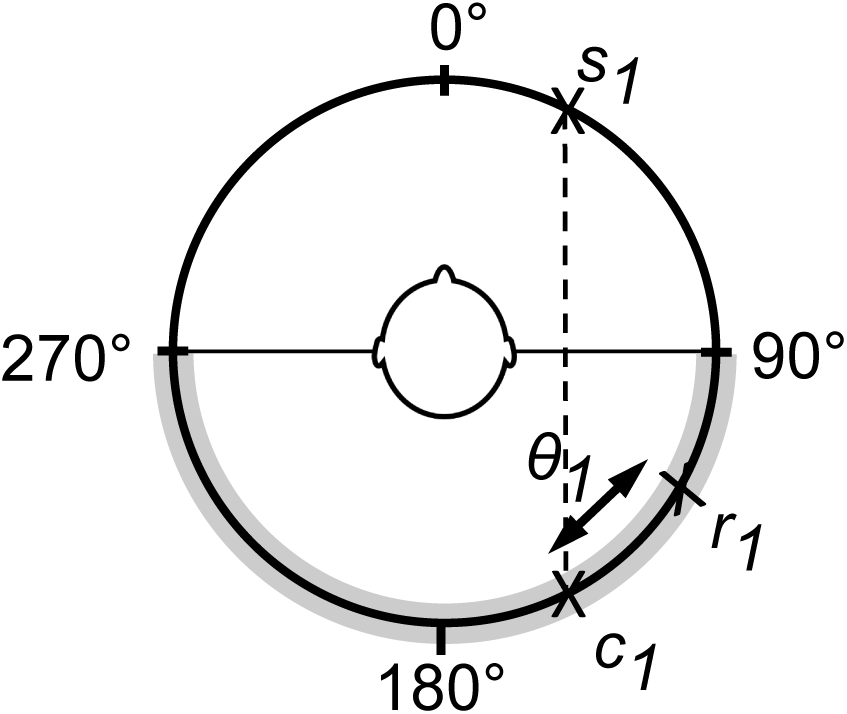
The FBC center c_1_ is the stimulus origin s_1_ mirrored on the interaural axis. The interaural axis corresponds to the line from 270° to 90°. Valid FBCs can occur within the light shaded stripe. The deviation of the response r_1_ from c_1_ is denoted with *θ*_1_. In this example, the response r_1_ lies in counter clockwise direction with respect to c_1_.

**Fig. 3.**
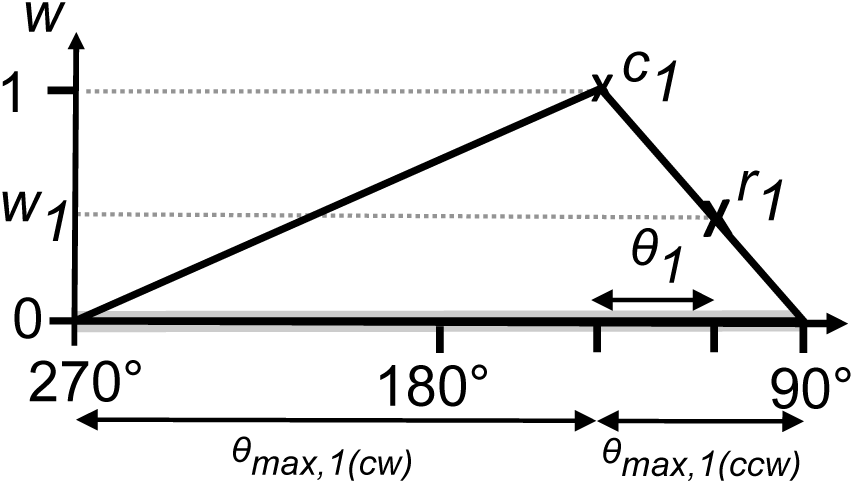
Weighting factor calculation for the example shown in Figure 2. The y-axis shows the front-back confusion weighting factor w and the x-axis shows the rear azimuth. The deviation of the response r_1_ from c_1_ is denoted by *θ*_1_. The response r_1_ lies in counter clockwise (ccw) direction with respect to s_1_, therefore *θ*_max,1_ is equal to *θ*_max,1_(ccw). The areas *θ*_max,1_(ccw) and *θ*_max,1_(cw) are limited by the interaural axis as defined in Figure 2.

To provide a subject-level FBC metric that considers deviations of the responses from the cone of confusion and their proximity to the interaural axis, we propose the FBC Score as defined in Equation 3.

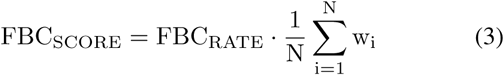

An FBC Score of 100% indicates that all responses were FBCs and exactly match the stimulus positions mirrored on the interaural axis. In contrast, an FBC Score of 0% would indicate that the responses did not contain any FBCs during the trial. For a calculation example with numeric data, a step-by-step guide for the calculation of the FBC Score with corresponding explanations is provided in the Appendix A. To facilitate the calculations, we implemented the procedure as a MATLAB function^1^. The script takes the sets of stimuli Ŝ and responses 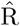 as input parameters. In addition to the calculations described above, a separate analysis of front-back and back-front confusions is performed.

## III. Results

For the CI users, the overall FBC Score is 27% in contrast to the almost twice as high FBC Rate of 47% (*p*< .001, two-sided Wilcoxon signed rank test). For the NH control group, the FBC Rate is 4 times as high as the FBC Score (FBC Score: 1%, FBC Rate: 4%; *p*= 0.062). Table I shows the results of the FBC Score analysis compared to the FBC Rate for each CI user and NH subject.

**TABLE I.** The table shows the results on subject-level for the front-back confusion (FBC) score analysis compared to the FBC Rate for each cochlear implant (CI) user and normal hearing (NH) subject.

| ID | FBC Rate (%) | FBC Score (%) |
| --- | --- | --- |
| CI01 | 60.0 | 41.3 |
| CI02 | 46.7 | 19.8 |
| CI03 | 53.3 | 28.8 |
| CI04 | 50.0 | 39.9 |
| CI05 | 46.7 | 22.7 |
| CI06 | 43.3 | 26.9 |
| CI07 | 40.0 | 20.1 |
| CI08 | 46.7 | 18.2 |
| CI09 | 33.3 | 15.4 |
| CI10 | 50.0 | 31.3 |
| CI11 | 40.0 | 23.4 |
| CI12 | 50.0 | 27.8 |
| NH01 | 0.0 | 0.0 |
| NH02 | 0.0 | 0.0 |
| NH03 | 0.0 | 0.0 |
| NH04 | 0.0 | 0.0 |
| NH05 | 10.0 | 4.2 |
| NH06 | 6.7 | 0.0 |
| NH07 | 16.7 | 9.4 |
| NH08 | 0.0 | 0.0 |
| NH09 | 6.7 | 1.3 |
| NH10 | 0.0 | 0.0 |
| NH11 | 0.0 | 0.0 |
| NH12 | 3.3 | 0.0 |

## IV. Conclusion and Discussion

For the evaluation of sound localization experiments, the errors are quantified in terms of their deviation from the stimulus position. However, for FBC analysis, thus far, only a rough distinction between “no FBC” and “FBC” is made, and the occurrences are counted. In this report, we propose a method for quantifying the severity of an FBC, allowing a more precise analysis of this phenomenon.

In the example presented here, the FBC Rate would indicate that the tested subjects are prone to FBCs (47%). However, the FBC Score shows that this assumption does not necessarily hold true since the impact of the FBCs is significantly smaller by a factor of 1.7 (27%). Therefore, the FBC Rate in our example includes many FBCs with a considerable deviation from the corresponding stimulus position on the cone of confusion, to which this phenomenon actually refers.

An alternative option to illustrate this influence on the phenomenon of FBCs would be to indicate, in addition to the FBC Rate, the localization error with respect to the stimulus’ position mirrored on the interaural axis. In our opinion, however, this approach may overestimate the importance of FBCs caused by stimuli near the interaural axis for two reasons: First, the influence of FBCs near the interaural axis is rather small in real life situations. Second, both error types, i.e., the FBC and localization errors, overlap for stimuli originating from close to the interaural axis, which can lead to distortions in the FBC analysis. By weighting the distance between the stimulus and the interaural axis and by considering the proximity of the stimulus to the interaural axis, this influence on the assessment of the FBC phenomenon can be mitigated. The effect of this weighting is particularly demonstrated by the 4 times lower FBC Score compared to the FBC Rate in the NH control group. Here, all FBC-causing stimuli originated from the measurement position with the smallest distance to the interaural axis.

The studies in [11], [12] defined responses as FBCs if the maximum deviation from the FBC center was less than or equal to a 40 or 45-degree angle. Such a static threshold value is not feasible in clinical studies with hearing-impaired subjects, due to the large differences in localization accuracy between the NH control and the test group [14]. An explanation of the underlying theoretical and psychoacoustic mechanism would be missing for the justification of such a threshold value. Besides the static threshold value, the iterative regression method applied in [11], [12] relies on the distribution and number of sound localization responses. The FBC Score does neither depend on a predefined threshold value nor on the quality or quantity of the localization results. This may be especially important when evaluating the performance of hearing-impaired patients in clinical trials. These localization results often show a wide individual variability [14] and consist of limited samples due to the limited testing time and subject attention.

## Supporting information

Matlab script for FBC calculations

## Appendix A

### Example calculation with numeric data

In the following, we illustrate the calculation of the FBC Score using a small example data set with 5 stimuli Ŝ and responses 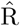 from a sound localization test in the horizontal plane (see Figure 4). Table II summarizes the results for the computational steps involved.

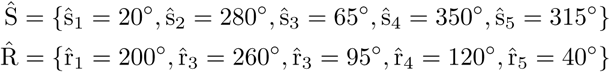

**TABLE II.** Data of the calculation example.

| Test item | Stimulus ($^{\circ}$) direction $s_i$ | Response ($^{\circ}$) direction $r_i$ | FBC center ($^{\circ}$) $c_i$ | Deviation ($^{\circ}$) $\theta_i$ | Maximum ($^{\circ}$) deviation $\theta_{\max,i}$ | Weighting factor $w_i$ |
| --- | --- | --- | --- | --- | --- | --- |
| 1 | 20 | 200 | 160 | 40 | 110 | 0.64 |
| 2 | 280 | 260 | 260 | 0 | 10 | 1.00 |
| 3 | 65 | 95 | 115 | 20 | 25 | 0.20 |
| 4 | 350 | 120 | 190 | 70 | 100 | 0.30 |

**Fig. 4.**
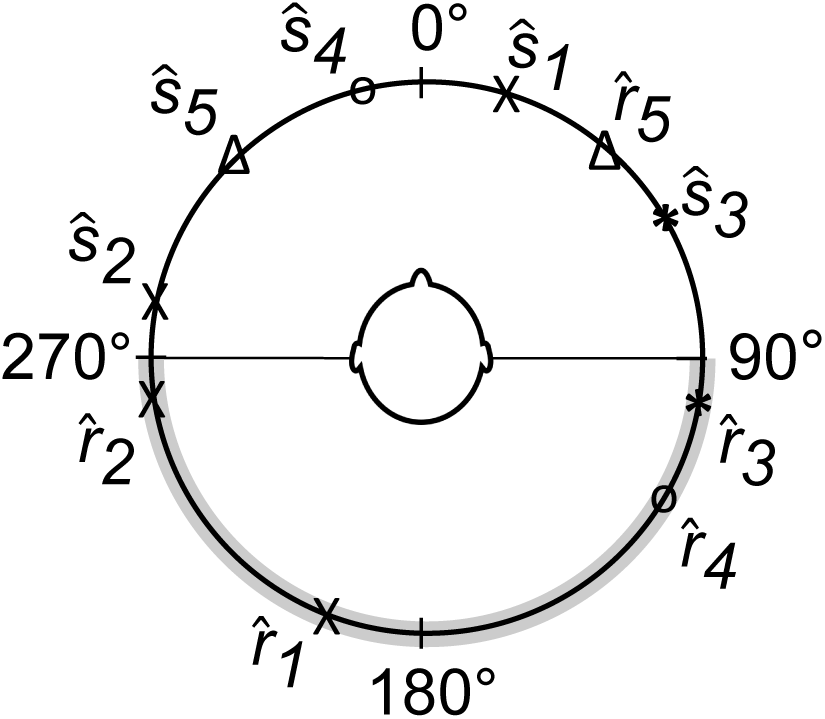
Visualization of the data for the example calculation of the front-back confusion (FBC) Score. Stimuli are indicated with Ŝ and responses with 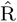. In this example, all stimuli lie inside the frontal azimuth and thus responses inside rear azimuth (light shaded area) are considered FBCs.

Step 1 - Calculate the FBC Rate:

First, the FBC Rate is calculated. It is defined as the rate of the number of responses crossing the interaural axis with respect to the number of presented stimuli (see Eq. 1). Only 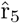 does not cross the interaural axis. Excluding ŝ_5_ and 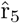 results in N = 4 pairs of stimuli s_i_ and responses r_i_ and an FBC Rate of 80%.

Step 2 - Calculation of c_i_ and *θ*_i_:

The FBC center c_i_ is obtained by mirroring the stimulus position s_i_ on the interaural axis. The deviation *θ*_i_ is calculated as the absolute difference between the response r_i_ and c_i_ measured over the minor arc. For example, for stimulus 3, we have *θ*_3_ = |r_3_ − c_3_| = |95° − 115°| = 20°.

Step 3 - Calculation of *θ*_max,i_:

The maximum deviation *θ*_max,i_ between r_i_and c_i_ is limited by the interaural axis. If r_i_ lies in clockwise direction of c_i_, the clockwise limit is applied (*θ*_max,i_= *θ*_max,i(cw)_). The same applies for the counter clockwise direction.

For example, with stimulus s_3_, the minor arc for the response r_3_ lies in counter clockwise direction to the FBC center c_3_, therefore the maximum deviation is *θ*_max,3_ = *θ*_max,3(ccw)_ = 25°.

Step 4 - Calculation of w_i_:

With *θ*_i_ and *θ*_max,i_ we compute the weighting factor w_i_ for each stimulus using equation (2).

Step 5 - FBC Score:

We now calculate the FBC Score as defined in Equation (3):

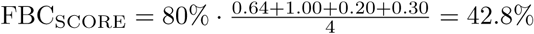

To facilitate the calculations, we implemented the procedure as a MATLAB function^1^.

## Acknowledgment

The authors would like to thank Christoph Schmid from the Department of ENT, Head and Neck Surgery, Bern University Hospital (Inselspital), for the recruitment of the CI users of this study.

## Footnotes

1 https://www.artorg.unibe.ch/research/hrl/data/fbc_score/

