## Supplementary material for "A Front-Back Confusion Metric in Horizontal Sound Localization: The FBC Score": Matlab script for FBC calculations: fbc_score.m

```
function [FBC_Analysis] = fbc_score(stimuli_raw_deg, responses_raw_deg)
clearvars -except stimuli_raw_deg responses_raw_deg
%--------------------------------------------------------------------------
%  This function calculates the FBC-Score: 
%  The FBC-Score introduces a weighting between actual FBCs and
%  normal localization errors. It allows to specify the influence of FBCs 
%  on a localization result in the azimuthal plane more precisely 
%  than with the FBC rate. 

%  FBC_Analysis.fbcs_and_bfcs.fbc_score_percent

%  If used, please cite:
%  ~~~~~~~~~~~~~~~~~~~~~~~~~~~~~~~~~~~~~~~~~~~~~~~~~~~~~~~~~~~~~~~~~~~~~~~~
%  Fischer, T., M. Caversaccio, W. Wimmer (2020). 
%  A Front-Back Confusion Metric in Horizontal Sound Localization: The FBC Score

%  or 

%  Pinna-Imitating Microphone Directionality Improves Sound Localization and Discrimination in Bilateral Cochlear Implant Users
%  Fischer T., Schmid S., Kompis M., Mantokoudis G., Caversaccio M., Wimmer W. (2020)
%  ~~~~~~~~~~~~~~~~~~~~~~~~~~~~~~~~~~~~~~~~~~~~~~~~~~~~~~~~~~~~~~~~~~~~~~~~
%
%   This code is part of the supplement material to the article:
%
%     Pinna-Imitating Microphone Directionality Improves Sound Localization and Discrimination in Bilateral Cochlear Implant Users
%     Fischer T., Schmid S., Kompis M., Mantokoudis G., Caversaccio M., Wimmer W. (2020)
%
%--------------------------------------------------------------------------
%% ASSIGNMENT OF TEXT DESCRIPTIONS TO VARIALBE NAMES:
% alpha_i:                      FBC-Area: Light shaded area in Figure 1, 
%                               corresponding to a FBC-Center c_i. Only 
%                               responses r_i that lie wihtin the 
%                               corresponding FBC-Area alpha_i, are 
%                               considered a FBC. The limits of the area 
%                               are defined by the interaural axis.
%--------------------------------------------------------------------------


%% FUNCTION PARAMETERS:
%--------------------------------------------------------------------------
% Input:
% stimuli_raw_deg:                      Vector with azimuths of the stimuli
% responses_raw_deg:                    Vector with azimuths of the responses

% Output:
% FBC_Analysis:                 		Struct containing FBC-Analysis

% Upper struct levels names explanations:
% ---------------------------------------    
% s_i_deg:              Stimuli causing a FBC response
% r_i_deg:              Responses which correspond to a FBC

% fbcs_and_bfcs:        FBCs and BFCs are treated equally
% only_bfcs:            Only BFCs are considered for caclations
% only_fbcs:            Only FBCs are considered for caclations
        
% Last struct level names explanations:
% --------------------------------------
% fbc_score_percent:            FBC-Score as introduced in (Fischer et al. 2020).
% number_of_fbcs:               Stimuli inside the FBC-Area alpha_i (= stimuli crossing the interaural axis (IA)).
% fbc_rate_percent:             Ratio of responses crossing the IA
%--------------------------------------------------------------------------

%--------------------------------------------------------------------------
%
%     FBC-Score Code (v1.0)
%     Copyright (C) 2020  Tim Fischer
%    .
%
%     This program is free software: you can redistribute it and/or modify
%     it under the terms of the GNU General Public License as published by
%     the Free Software Foundation, either version 3 of the License, or (at
%     your option) any later version.
%
%     This program is distributed in the hope that it will be useful, but
%     WITHOUT ANY WARRANTY; without even the implied warranty of
%     MERCHANTABILITY or FITNESS FOR A PARTICULAR PURPOSE.  See the GNU
%     General Public License for more details.
%
%     You should have received a copy of the GNU General Public License
%     along with this program.  If not, see <http://www.gnu.org/licenses/>.
%
%--------------------------------------------------------------------------
%% INPUT PARAMETERS:
%--------------------------------------------------------------------------
% These are the stimuli and responses from the example in the publication
if (nargin < 2)
    stimuli_raw_deg                 =   [20  280 65 350 315]';
    responses_raw_deg               = 	[200 260 95 120 40]';
end
if (length (stimuli_raw_deg) ~= length(responses_raw_deg))
    error('The number of stimuli and responses must be equal');
end
% Make sure stimuli and responses are correctly formatted (0� to 360�)
stimuli_raw_deg                     =   round(mod(stimuli_raw_deg, 360), 1);
responses_raw_deg                   =   round(mod(responses_raw_deg, 360), 1); 
% Store stimuli and responses
FBC_Analysis.stimuli_raw            =   stimuli_raw_deg;
FBC_Analysis.responses_raw          =   responses_raw_deg;


%% DEFINITION OF CONSTANTS:
%--------------------------------------------------------------------------
% Angle defintions
FRONT_DEG                           =   0;
RIGHT_SIDE_DEG                      =   90;
BACK_DEG                            =   180;
LEFT_SIDE_DEG                       =   270;
FBC_Analysis.FRONT_DEG              =   FRONT_DEG;
FBC_Analysis.BACK_DEG               =   BACK_DEG;
FBC_Analysis.LEFT_SIDE_DEG          =   LEFT_SIDE_DEG;
FBC_Analysis.RIGHT_SIDE_DEG         =   RIGHT_SIDE_DEG;


% Defintion of analysis parameters
CONFIGURATIONS = {'MERGED_FBC_AND_BFC_ANALYSIS', 'ONLY_BFC_ANALYSIS', 'ONLY_FBC_ANALYSIS'};


%% ITERATE OVER EACH SETTING:
%--------------------------------------------------------------------------
% Volatile analysis parameters
idx                                 =   1;

while idx <= length(CONFIGURATIONS)
    tmp_cnfg    =   char(CONFIGURATIONS(idx));
    idx         =   idx + 1;
    switch tmp_cnfg
        case 'MERGED_FBC_AND_BFC_ANALYSIS' 
            % This case covers the analysis as described in the paper.
            is_merged_fbc_and_bfc_analysis  =   true;
            is_only_bfc_analysis            =   false;
            is_only_fbc_analysis            =   false;
        case 'ONLY_BFC_ANALYSIS'
            is_merged_fbc_and_bfc_analysis  =   false;
            is_only_bfc_analysis            =   true;
            is_only_fbc_analysis            =   false;
        case 'ONLY_FBC_ANALYSIS'
            is_merged_fbc_and_bfc_analysis  =   false;
            is_only_bfc_analysis            =   false;
            is_only_fbc_analysis            =   true;
        otherwise
            error('Unexpected cnfg type. No FBC analysis performed.')
    end 
    
    
    %% EXCLUDING NON FBC STIMULI AND RESPONSES:
    %----------------------------------------------------------------------
    % Exclude stimuli and responses at the interaural axis. 
    % They also do not count for the fbc rate.
    right_side_stimuli_idcs = find(stimuli_raw_deg == RIGHT_SIDE_DEG);
    left_side_stimuli_idcs = find(stimuli_raw_deg == LEFT_SIDE_DEG);
    right_side_responses_idcs = find(responses_raw_deg == RIGHT_SIDE_DEG);
    left_side_responses_idcs = find(responses_raw_deg == LEFT_SIDE_DEG);
    exclude_side_stimuli_idcs = [right_side_stimuli_idcs; left_side_stimuli_idcs; right_side_responses_idcs; left_side_responses_idcs];
    stimuli_raw_deg(exclude_side_stimuli_idcs) = [];
    responses_raw_deg(exclude_side_stimuli_idcs) = [];
    
    [s_i_deg, r_i_deg] = get_r_i_crossing_the_IA(stimuli_raw_deg, responses_raw_deg);

    
    % Exclude stimuli in the frontal/dorsal azimuth positions, with
    % corresponding responses, from the FBC analysis (if desired by the
    % setting)
    if ( is_only_bfc_analysis == true )
        idcs_frontal_stimuli            =   find(get_is_azimuth_frontal( s_i_deg ));
        s_i_deg(idcs_frontal_stimuli)   =   [];
        r_i_deg(idcs_frontal_stimuli)   =   [];
    elseif (is_only_fbc_analysis == true)
        idcs_dorsal_stimuli             =   find(~get_is_azimuth_frontal( s_i_deg ));
        s_i_deg(idcs_dorsal_stimuli)    =   [];
        r_i_deg(idcs_dorsal_stimuli)    =   [];
    end
    
    %% CALCULATE THE FBC-CENTERS c_i AND FBC-AREA alpha_i
    %----------------------------------------------------------------------
    % FBC-Centers:
    c_i                               =   get_fbc_centers(s_i_deg);
    
    % FBC-Areas:
    alpha_i                           =   get_fbc_areas(c_i);


    %% CALCULATE THE OFFSET OF THE FBC-CENTER (=theta_i) 
    % AND THE MAXIMUM POSSIBLE OFFSET theta_{max,i}
    % ---------------------------------------------------------------------
    % Size of minor arcs between response r_i and c_i) (=theta_i):
    theta_i                     =   get_minor_arc(c_i,r_i_deg);
    % Maximum possible deviation in cw or ccw direction from c_i 
    % to be still counted as FBC (limits are defined by the interaural axis)
    theta_max_i                 =   get_theta_max_size(c_i, r_i_deg, alpha_i);

    
    %% CALCULATE THE FBC-WEIGHTING w_i
    % ---------------------------------------------------------------------
    % Ratio of theta_max_i w.r.t. cw or ccw interval limit of alpha_i 
    w_i                       =   1 - round((theta_i ./ theta_max_i), 2);
    if ( any(w_i > 1) || any(w_i < 0) )
        error('Weighting is outside range')
    end
    %% CALCULATE THE FBC-RATE
    %----------------------------------------------------------------------    
    % FBC DEFINITION, WHERE ALL CROSSINGS OF THE IA ARE COUNTED AS FBC
    % Number of FBCs and FBC-Rate (Responses crossing the interaural axis,
    % no further conditions need to be met to be counted as fbc)
    number_of_fbcs              =   length(r_i_deg);
    fbc_rate_percent            =   round((number_of_fbcs/length(responses_raw_deg)) * 100, 1);
    if (isnan(fbc_rate_percent))
        fbc_rate_percent = 0.0;
    end
    
    %% CALCULATE THE FBC-SCORE
    %----------------------------------------------------------------------
    fbc_score_percent               =   round(fbc_rate_percent*mean(w_i), 1);
    if (isnan(fbc_score_percent))
        fbc_score_percent = 0.0;
    end
    if (fbc_score_percent > fbc_rate_percent)
        error('The FBC score can never be higher than the FBC rate');
    end
    
    %% OUTPUT :
    %----------------------------------------------------------------------
    % Store all FBC-Metrics with corresponding setting in the FBC-Analysis
    % struct
    if (is_merged_fbc_and_bfc_analysis)
        % FB an BF confusions are treated equally
        FBC_Analysis.fbcs_and_bfcs.fbc_score_percent         =   fbc_score_percent;
        FBC_Analysis.fbcs_and_bfcs.number_of_fbcs            =   number_of_fbcs;
        FBC_Analysis.fbcs_and_bfcs.fbc_rate_percent          =   fbc_rate_percent;
        FBC_Analysis.stimuli_deg                             =   s_i_deg;
        FBC_Analysis.responses_deg                           =   r_i_deg;
    elseif(is_only_bfc_analysis)
        % Only back-front confusions are analyzed
        FBC_Analysis.only_bfcs.fbc_score_percent             =   fbc_score_percent;
        FBC_Analysis.only_bfcs.number_of_fbcs                =   number_of_fbcs;
        FBC_Analysis.only_bfcs.fbc_rate_percent              =   fbc_rate_percent;    
    elseif(is_only_fbc_analysis)
        % Only front-back confusions are analyzed
        FBC_Analysis.only_fbcs.fbc_score_percent             =   fbc_score_percent;
        FBC_Analysis.only_fbcs.number_of_fbcs                =   number_of_fbcs;
        FBC_Analysis.only_fbcs.fbc_rate_percent              =   fbc_rate_percent;               
    end

end
% Checksum for correctness of the values
checksum_fbc_analysis = FBC_Analysis.only_fbcs.number_of_fbcs + FBC_Analysis.only_bfcs.number_of_fbcs;
if ( checksum_fbc_analysis ~= FBC_Analysis.fbcs_and_bfcs.number_of_fbcs )
    error('There is something wrong. fbcs+bfcs != (fbcs and bfcs)');
end


    %--------------------------------------------------------------------------
    %--------------------------------------------------------------------------
    %% NESTED FUNCTIONS (HELPER FUNCTIONS):
    %--------------------------------------------------------------------------
    %--------------------------------------------------------------------------
    function [minor_arcs]       = get_minor_arc(angle1_deg,angle2_deg)
    %  This function calculates the minor arc between two angles on the
    %  circle

    %% PARAMETERS:
    %--------------------------------------------------------------------------
    % Input:
    % angle1_deg:              Vector with azimuths 
    % angle2_deg:              Vector with azimuths

    % Output:
    % minor_arcs:              Vector returning the minor arcs   
    minor_arcs = abs(rad2deg(angle(exp(deg2rad(angle1_deg)*1i)./exp(deg2rad(angle2_deg)*1i))));        
    end
    function [theta_max_i_out] = get_theta_max_size(c_i_, r_i_deg_, alpha_i_)
    %  This function calculates the maximum size of the minor arc between the 
    %  response r_i and the FBC-Center c_i (=theta_max_i). The size of the arc
    %  defines the relative punishment of r_i_s being away from c_i.

    %% PARAMETERS:
    %--------------------------------------------------------------------------
    % Input:
    % c_i_:             Vector with FBC-Center azimuths
    % r_i_deg_:         Vector with azimuths of the responses
    % alpha_i_:         Start- and end azimuths (cw direction) of the FBC-Areas

    % Output:
    % theta_max_i_out:  Size of the max. possible deviation of r_i to
    %                   c_i_

    %% DEFINITION OF CONSTANTS:
    %--------------------------------------------------------------------------
    CCW_LIMIT       =  1;
    CW_LIMIT        =  2;

    %% THETA_MAX_I CALCULATION
    %--------------------------------------------------------------------------
    theta_max_i_out                         =   nan(length(r_i_deg_),1);
    for i = 1 : length(r_i_deg_)
        tmp_response                        =   r_i_deg_(i);
        tmp_c_i_                            =   c_i_(i);
        % Response cw or ccw of FBC-Center (0:false, 1:true)
        tmp_distance    =   get_minor_arc(tmp_c_i_,tmp_response);
        if (round(mod(tmp_c_i_+tmp_distance,360),1) == tmp_response)
            is_response_in_ccw_direction    =   false;
        else
            is_response_in_ccw_direction    =   true;
        end
        % Take the corresponding limit max\alpha_{i,cw(ccw)} and calculate the
        % size of theta_max_i_out
        if (is_response_in_ccw_direction == true)
            tmp_alpha_max                   =   alpha_i_(i,CCW_LIMIT);
        else
            tmp_alpha_max                   =   alpha_i_(i,CW_LIMIT);
        end
        theta_max_i_out(i) = round(get_minor_arc(tmp_c_i_,tmp_alpha_max),1);
    end
    if ( any(theta_max_i_out) > 180)
        error ('Max deviation cannot be higher than 180 degree');
    end

    end


    function [s_i_crossing_IA_out, r_i_crossing_IA_out] = get_r_i_crossing_the_IA(s_i_deg_, r_i_deg_)
    %  This function the responses (with corresponding stimuli) which cross the 
    %  interaural axis (IA) w.r.t. the azimuth of the stimulus

    %% PARAMETERS:
    %--------------------------------------------------------------------------
    % Input:
    % s_i_deg_:              Vector with azimuths of the stimuli
    % r_i_deg_:              Vector with azimuths of the responses

    % Output:
    % r_i_crossing_IA_out:   Vector returning the responses, crossing the IA

    r_i_crossing_IA_out = [];
    s_i_crossing_IA_out= [];
    for i = 1:length(s_i_deg_)
        tmp_stimulus_deg                =   s_i_deg_(i);
        tmp_response_deg                =   r_i_deg_(i);
        is_stimulus_in_frontal_sphere   =   get_is_azimuth_frontal(tmp_stimulus_deg);
        is_response_in_frontal_sphere   =   get_is_azimuth_frontal(tmp_response_deg);

        % Stimulus and response are not in the the same sphere
        if (sum([is_stimulus_in_frontal_sphere,is_response_in_frontal_sphere]) == 1)
            r_i_crossing_IA_out(end+1,1)  =   tmp_response_deg;
            s_i_crossing_IA_out(end+1,1)  =   tmp_stimulus_deg;
        end
    end
    end


    function [alpha_i_out] = get_fbc_areas(c_i_)
    %  This function calculates the FBC-Areas alpha_i. (Only responses that lie
    %  wihtin the corresponding FBC-Area alpha_i, are considered a FBC)

    %% PARAMETERS:
    %--------------------------------------------------------------------------
    % Input:
    % c_i_:          Vector with FBC-Center azimuths

    %% DEFINITION OF CONSTANTS:
    %--------------------------------------------------------------------------
    CCW_LIMIT           =   1;
    CW_LIMIT            =   2;
    
    % Output:
    % alpha_i_out:   Start- and end azimuths (cw direction) of the FBC-Areas               

    %% Area of MAX_ALPHA_SIZE_DEG around the FBC-Center c_i (minor
    % arc). The area is defined by c_i+theta_{max,i(cw)} and c_i-theta_{max,i(ccw)}
    % and thus is restricted by the interaural axis (see Figure 2).
    %--------------------------------------------------------------------------
    alpha_i_out     =   nan(length(c_i_),2); 
    for i = 1 : length(c_i_)        
        if (get_is_azimuth_frontal(c_i_(i)) == true)
            alpha_i_out(i,CCW_LIMIT) = LEFT_SIDE_DEG;
            alpha_i_out(i,CW_LIMIT)  = RIGHT_SIDE_DEG; 
        else
            alpha_i_out(i,CCW_LIMIT) = RIGHT_SIDE_DEG;
            alpha_i_out(i,CW_LIMIT)  = LEFT_SIDE_DEG; 
        end
    end
    end


    function [azimuth_is_frontal_out] = get_is_azimuth_frontal(azimuth_deg_)
    %  This function returns true, if the azimuth_deg is within the frontal
    %  sphere of a circle (>270� && < 90�).

    %% PARAMETERS:
    %--------------------------------------------------------------------------
    % Input:
    % azimuth_deg_:             Vector with azimuths in deg

    % Output:
    % azimuth_is_frontal_out:   Vector returning true, if azimuth is in frontal
    %                           sphere of the circle

    %% IS IN FRONTAL SPHERE CHECK:
    %--------------------------------------------------------------------------
    azimuth_is_frontal_out              =   nan(length(azimuth_deg_), 1);
    for i = 1 : length(azimuth_deg_)
        tmp_azimuth_deg_                =   azimuth_deg_(i);
        tmp_azimuth_is_frontal_out      =   find((tmp_azimuth_deg_ >= LEFT_SIDE_DEG) || (tmp_azimuth_deg_ <= RIGHT_SIDE_DEG));
        if (isempty(tmp_azimuth_is_frontal_out))
            tmp_azimuth_is_frontal_out  =    false;
        else
            tmp_azimuth_is_frontal_out  =    true;
        end
        azimuth_is_frontal_out(i)       =   tmp_azimuth_is_frontal_out;
    end
    end


    function [c_i_out] = get_fbc_centers( s_i_deg_ )
    %  This function calculates the FBC-Centers c_i. 
    %  FBC-Centers c_i are the stimulis positions mirrored at the interaural-
    %  axis (IA).

    %% PARAMETERS:
    %--------------------------------------------------------------------------
    % Input:
    % s_i_deg_:      Vector with azimuths of the stimuli

    % Output:
    % c_i_out:       Vector with FBC-Center azimuths

    %% FBC-Center calculation
    %--------------------------------------------------------------------------
    c_i_out = [];
    for i = 1:length(s_i_deg_)
        tmp_stimuli_deg                 =   s_i_deg_(i);

        is_stimulus_in_right_sphere     =   find ((tmp_stimuli_deg >= FRONT_DEG) & (tmp_stimuli_deg <= BACK_DEG));
        is_stimulus_in_left_sphere      =   find ((tmp_stimuli_deg >= BACK_DEG));

        if (is_stimulus_in_right_sphere)
            distance_to_IA      =   RIGHT_SIDE_DEG - tmp_stimuli_deg;
            % Stimulus is in the frontal azimuth position
            c_i_out(end+1)      =   RIGHT_SIDE_DEG + distance_to_IA;
        elseif (is_stimulus_in_left_sphere)
            distance_to_IA      =   LEFT_SIDE_DEG - tmp_stimuli_deg;
            c_i_out(end+1)      =   LEFT_SIDE_DEG + distance_to_IA;
        end
    end

    if (length(c_i_out) ~= length(s_i_deg_))
        error('Not every stimuli has been assigned to a FBC-Center');
    end
    if (isempty(c_i_out))
        c_i_out = double.empty(0,1);
    else
        c_i_out = round(c_i_out',1);
    end
    end
end    
    
```
